# Functional ultrasound imaging of recent and remote memory recall in the associative fear neural network in mice

**DOI:** 10.1101/2021.11.13.468469

**Authors:** Gillian Grohs-Metz, Rebecca Smausz, John Gigg, Tobias Boeckers, Bastian Hengerer

## Abstract

Emotional learning and memory are affected in numerous psychiatric disorders. At a systems level, however, the underlying neural circuitry is not well defined. Rodent fear conditioning (FC) provides a translational model to study the networks underlying associative memory retrieval. In the current study, functional connectivity among regions related to the cue associative fear network were investigated using functional ultrasound (fUS), a novel imaging technique with great potential for detecting regional neural activity through cerebral blood flow. Behavioral fear expression and fUS imaging were performed one and thirty-one days after FC to assess recent and remote memory recall. Cue-evoked increases in functional connectivity were detected throughout the amygdala, with the lateral (LA) and central (CeA) amygdalar nuclei emerging as major hubs of connectivity, though CeA connectivity was reduced during remote recall. The hippocampus and sensory cortical regions displayed heightened connectivity with the LA during remote recall, whereas interconnectivity between the primary auditory cortex and temporal association areas was reduced. Subregions of the prefrontal cortex exhibited variable connectivity changes, where prelimbic connectivity with the amygdala was refined while specific connections between the infralimbic cortex and amygdalar subregions emerged during remote memory retrieval. Moreover, freezing behavior positively correlated with functional connectivity between hubs of the associative fear network, suggesting that emotional response intensity reflected the strength of the cue-evoked functional network. Overall, our data provide evidence of the functionality of fUS imaging to investigate the neural dynamics of memory encoding and retrieval, applicable in the development of innovative treatments for affective disorders.

**Highlights:** Functional ultrasound imaging can elucidate fear associated neural networks

Freezing behavior correlates with cue-evoked functional connectivity changes

The lateral and central amygdalar nuclei are major hubs in the fear network

The hippocampus is active during recent and remote cued fear memory retrieval

Connectivity profiles of the prelimbic and infralimbic areas vary in remote recall

## 1. Introduction

Learning and memory are of major importance for an organism’s survival, enabling it to adapt to the challenges of our complex, ever-changing environment. There is accumulating evidence that memories are stored in a distributed neuronal ensemble, or memory engram [1,2]. Underlying a memory are stages of encoding, consolidation, and retrieval where the engram is thought to become permanently encoded through molecular processes such as synaptic plasticity and strengthening [1]. During engram reactivation, a memory trace can be detected through the activation of specific neural networks, visualized through changes in functional connectivity. The capacity for dynamic and adaptable learning is attenuated in many psychiatric disorders [3]. Therefore, elucidating the neural circuitry underlying these processes and how they are affected in disease is crucial for developing strategies to restore and maintain these cognitive abilities. More specifically, studying emotional learning is of relevance for affective and anxiety-related conditions.

Fear conditioning (FC) is one of the simplest and best characterized models to study the neural networks underlying emotional learning in the laboratory setting. It builds upon the classical Pavlovian conditioning principle, namely teaching an animal model, most often a rodent, to associate an emotionally relevant stimulus, called the unconditioned stimulus (US), with a naturally neutral one, called the conditioned stimulus (CS). Thus, the US is often a mild foot-shock, which is innately aversive and causes avoidance behavior, while the CS could be, for example, a tone, to which the animals initially show no response. However, after learning that the tone predicts the shock, animals will exhibit an emotional response upon exposure to the CS alone; thus, an associative memory of the CS-US pairing is formed [4]. There is extensive literature on this associative learning model, given how simple, cost-effective and quickly-acquired it is, as a single training session is sufficient for inducing the formation of a lasting and robust memory [5–7].

Although FC has been used for decades to model associative learning, until recently there has been little consensus about the brain regions involved in fear memory formation and long-term storage and retrieval [8]. Early studies indicated that certain brain regions can be pinpointed as the seat of fear memory storage; these include the basolateral complex of the amygdala, the emotion processing center of the brain, which can be sub-divided into the lateral (LA), basolateral (BLA) and basomedial (BMA) amygdalar nuclei [5,6]. Indeed, lesions of the basolateral complex abolish fear memory recall, regardless of whether being made before or after conditioning [6]. However, more recent studies point out that the amygdala is necessary, but not sufficient for permanent fear memory storage [8], which is rather encoded in distributed neural circuits [9,10]. Current evidence suggests that these include the amygdala, with both the basolateral complex and central nucleus (CeA) [5,11]; the sensory cortices, namely the auditory [8] and somatosensory areas [12]; and the prefrontal cortex (PFC), divided into the prelimbic (PL) and infralimbic (IL) regions [13,14]. While the role of the hippocampus (HC) is well established in contextual fear memory [15], its role in cued fear memory storage is still controversial. A number of studies have found that hippocampal involvement ceases after memory encoding, whereas others have found evidence that the hippocampus maintains a role in the retrieval of the engram through processes such as recontextualization and hippocampal indexing [16–19].

The distributed nature of fear representation strongly supports the use of circuit level analytical techniques to investigate the fear memory trace. New imaging techniques such as functional ultrasound (fUS) enable non-invasive and repeatable network-wide coverage with an exceptionally high signal to noise ratio [20–22]. By utilizing plane wave, ultrafast Doppler technology, fUS imaging can detect dynamic changes in local vasculature at a higher spatial and temporal resolution than those achieved with positron emission tomography (PET) and functional magnetic resonance imaging (fMRI) [23–25]. The temporal correlation of low frequency hemodynamic oscillations from a priori identified regions of interest (ROIs), a measure of functional connectivity, can be used to investigate network dynamics [21,26,27]. These neural network signatures are also maintained under sedation and light anesthesia. Indeed, robust resting state network analysis and stimulus-evoked functional hyperemia have been detected in anesthetized rodents [21,26,28]. Light sedation such as that achieved with dexmedetomidine, a potent α_2_ adrenergic receptor agonist, has been found to preserve functional connectivity and neural responsiveness to stimuli [26,29]. The silent and non-invasive nature of fUS imaging, in combination with the sedation provided by dexmedetomidine, provide an ideal set up for an auditory associated memory retrieval test. Thus, the present study aimed to investigate circuit wide cue-evoked functional connectivity changes following an associative fear learning task in mice using the technological advantages of fUS. In doing so, we sought to investigate the neural circuitry underlying emotional memory formation and long-term recall and to correlate these measures with behavioral expression.

## 2. Materials and Methods

### 2.1. Animals and license

All experiments were approved by the appropriate authority (Regierungspräsidium Tübingen, Germany) in accordance with European Directive 2010/63/EU and performed in an AAALAC (Association for Assessment and Accreditation of Laboratory Animal Care International) accredited facility. Experiments were performed on C57BL/6J male mice (Charles River Research Models and Services Germany GmbH, 7-8 weeks old, 21-23g at study onset) maintained on a normal 12:12 h light-dark cycle (lights on at 6 am). Mice were housed individually to eliminate aggression and injury and given *ad-libitum* access to normal mouse chow and water. After one week of acclimation, mice were habituated to handling for 10-15 minutes on three consecutive days prior to the beginning of the study in both the fUS and fear conditioning experimental rooms. All measurements were performed during light-phase.

### 2.2. Experimental Timeline

The fUS imaging sessions were run from 7 am to 1 pm with three mice per day. A baseline measurement was performed one day before fear conditioning (Day -1), followed by imaging sessions 2 and 31 days after task acquisition (Fig. 1). Fear conditioning (FC) was carried out at 2 pm on the day following baseline imaging (Day 0) with three mice simultaneously: 1 mouse per FC box. Recent recall of fear expression (FE) to tone was evaluated 24 h after conditioning, and FE to the conditioning context was carried out at 6 pm on the same day (Day 1). An imaging session to evaluate recent recall was performed the next day (Day 2). Remote recall was investigated via imaging after 31 days and behaviorally after 32 days (Fig. 1).

**Fig. 1.**
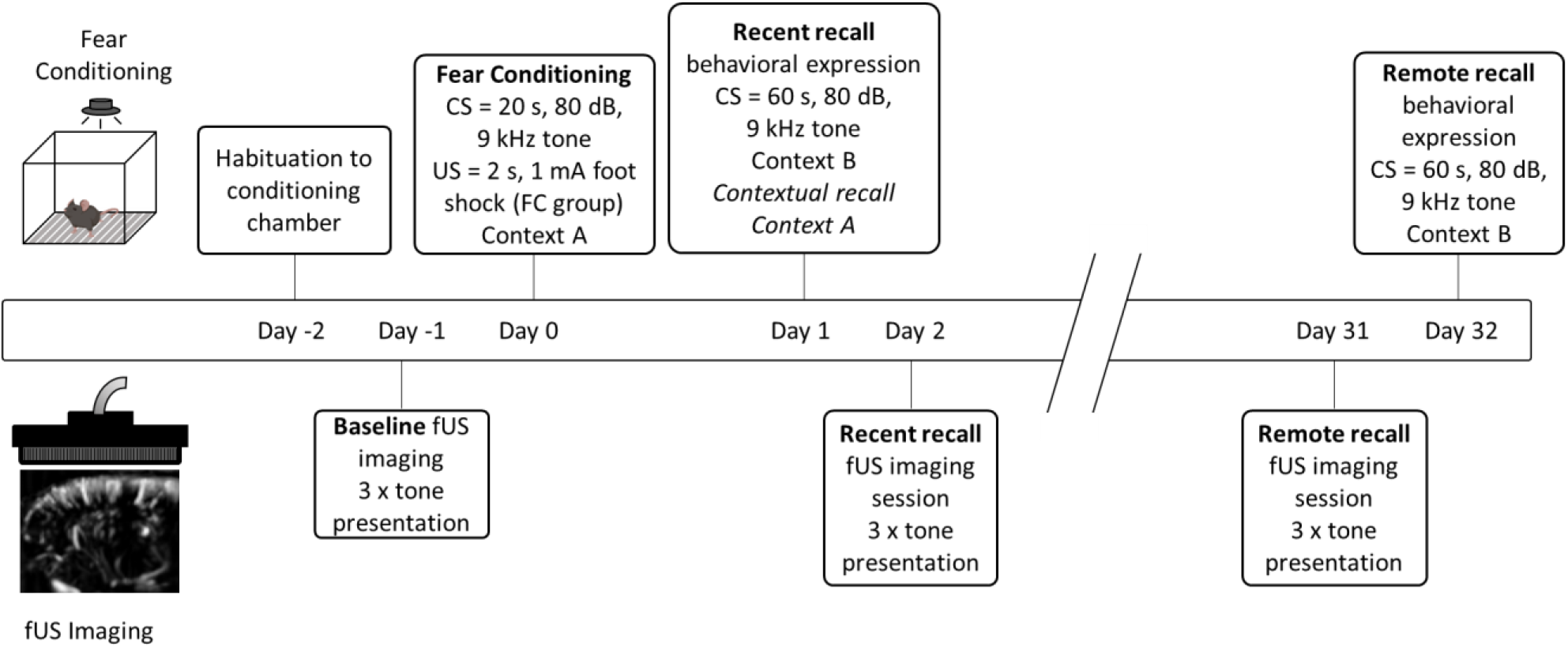
Training and imaging timeline. FC animals were exposed to a single tone - foot shock pairing while FT animals were only exposed to the tone. Recent and remote behavioral and functional connectivity responses were tested after 1 and 31 days.

### 2.3. Fear conditioning and behavioral expression

Behavioral experiments were carried out in four identical chambers (VFC-008, Med Associates, Vermont, USA), each enclosed in a sound attenuating box (NIR-022SD, Med Associates, Vermont, USA). VideoFreeze SOF-843 software (MedAssociates, Vermont, USA) was used to create protocols, run experiments and extract freezing data (detailed in [30]). In brief, each chamber consisted of a stainless-steel grid floor through which the electric shock (US) was delivered, a speaker in the side wall for tone (CS) delivery, and a fan for air circulation and constant background noise (65 dB) generation. An overhead LED-based light source provided white and near-infrared light, the latter being used by a VID-CAM-MONO-5 USB video camera to detect motion and freezing. Freezing was defined as the absence of all movement apart from that related to breathing. The minimum freezing duration was taken as 1 second in which the detected motion fell below the threshold (Motion Index Threshold = 18) [30]. Sessions were recorded at 30 frames per second. Mice were habituated to the chamber and grid floor for 15 minutes on the day prior to beginning the study (Day -2, Fig. 1) and were transferred in their home cages to the testing room 30 minutes prior to experiments. The FC and FE protocols were adapted from published literature [31,32] as follows: during FC, each mouse was placed in a separate chamber, which was lightly wiped with 75 % ethanol prior to start. The house light was set to 35 lux, referred to as ‘context A’. After three minutes of free exploration, a 20-second continuous tone (80 dB, 9 kHz, 10 ms rising and falling time) was played and co-terminated with a 2-second, 1 mA electric shock delivered through the grids, followed by one minute of free exploration before returning to the home cage. Mice were then left undisturbed in their home cages for 24 hours, after which FE to the CS was carried out in a novel context, referred to as ‘context B’. This consisted of completely covering the grids with a smooth flat white acrylic sheet, on top of which bedding was added to stimulate motility. One black and one grey acrylic sheet covered the left and right walls, respectively, and the house-light was increased to 50 lux. Mice freely explored the novel context for 3 min, then the same 80 dB, 9 kHz tone was played for 1 min (3 × 20 s bouts with 1 s in between), but without a shock, and again animals were returned to their home cages 1 min after the tone ended. The specificity of the freezing response to tone was tested on the same afternoon by re-exposing the animals for 5 min to context A without delivering the tone or the shock, to evaluate the contextual freezing response. Control animals were only exposed to the unconditioned tone (UT) without shock (n = 12), all other handling and experimental conditions were identical to the fear conditioned (FC) group (n = 12).

### 2.4. fUS imaging

Sedation was induced with 5 % isoflurane in a chamber and maintained at 1 % upon fixation in a stereotaxic frame (David Kopf Instruments, USA) positioned on an anti-vibration table (CleanBench™, TMC, Germany). Core temperature was maintained at 37 °C with a homeothermic warming system (PhysioSuite, Kent Scientific Corporation, USA). Meloxicam (Metacam®, Boehringer Ingelheim, DE) was given subcutaneously at 0.05 mg/kg to mitigate discomfort from stereotactic fixation. Non-rupture ear bars were placed ∼1 mm anterior to the ear canals with sufficient pressure to keep the head secure without causing tissue damage to allow full acoustic sensation by the mouse. After close electrical shaving of the hair on the scalp, a catheter was inserted into the dorsal side of the neck and a bolus of 0.067 mg/kg dexmedetomidine hydrochloride (Tocris, USA; freshly prepared and filtered) in physiological saline was administered, directly followed by continuous infusion at 0.2 mg/kg/h, 5 mL/kg flow. Ten minutes into infusion, gradual isoflurane turn-down was started at 0.1% per minute with total cessation 20 minutes into infusion. Functional imaging was initiated no earlier than 10 minutes after isoflurane termination.

Dedicated small animal fUS hardware and software (Iconeus, Paris, France) were used for all imaging sessions. A linear ultrasonic probe (15 MHz central frequency, Iconeus, Paris, France) with 128 piezoelectric transducers connected to an ultrafast ultrasound scanner (Iconeus One - 128 channels, Iconeus, Paris, France) was used to emit ultrasonic plane waves and to receive and process backscattered Power Doppler signal. fUS image compilation has been described in depth previously [20,21,26,28,33], and results in an image compiled from 200 compounded ultrafast frames acquired at a 500 Hz frame rate, each of those being the sum of ultrasonic echoes acquired from a set of 11 tilted plane waves (-10 to 10°, separated by 2°) emitted at a 5500 Hz pulse repetition frequency. The final sequence of functional images yielded a 2.5 Hz frame rate (1 power Doppler image acquired every 400 ms) with a spatial resolution of 100 × 100 × 400 μm. After animal fixation, scalp preparation and infusion, the scalp was covered with an isotonic coupling gel and the ultrasonic probe was lowered to 1 mm from the scalp for complete immersion into the gel. Probe position was calculated via the Iconeus Brain Positioning System [34] over an oblique plane that enabled imaging coverage of the majority of neuroanatomical regions of interest (ROIs) linked with fear networks (Fig. 2) [9,10]. Power Doppler images were acquired for 45 minutes where three 62 s tone blocks (3 × 20 s continuous 80 dB, 9 kHz tones with 1 s interval between) were presented via a mini loudspeaker placed near the animal’s head. The first tone block was initiated after 15 minutes and the subsequent tone blocks at 10-minute intervals to ensure a return to baseline neural connectivity. At the end of each imaging session, dexmedetomidine was antagonized with subcutaneous atipamezole (Alzane, Zoetis, Germany) at a dose of five times of total administered dexmedetomidine. Animals typically showed full reversal of sedation 10 minutes after atipamezole administration.

**Fig. 2.**
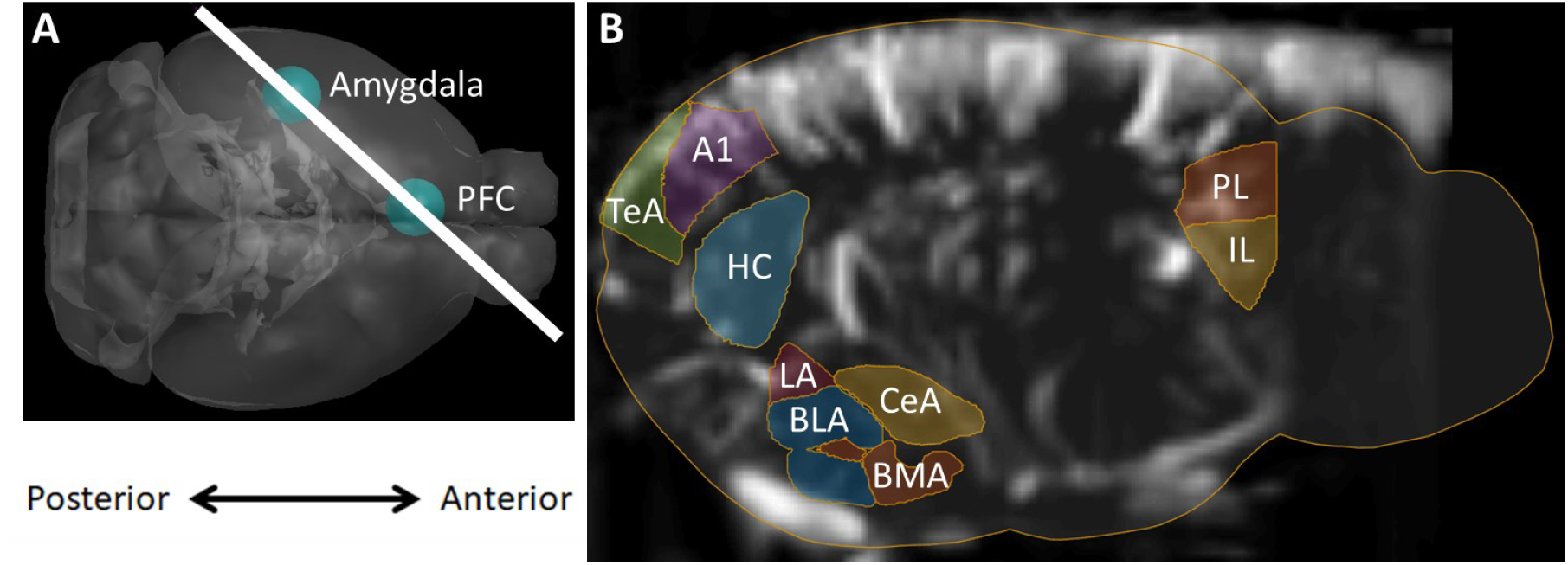
Oblique fUS imaging plane covering the ROIs in the left hemipshere. (A) 3D registration enabled linear probe positioning (white bar) through the PFC and amygdala (cyan spheres). (B) Anatomical delineations derived from the Allen Common Coordinate Framework overlaid on a representative fUS Doppler image show coverage of ROIs. PL prelimbic cortex, IL infralimbic cortex, A1 primary auditory cortex, TeA temporal association area, HC hippocampus, LA lateral amygdala, BLA basolateral amygdala, BMA basomedial amygdala, CeA central amygdalar nucleus.

### 2.5. Data Processing and Statistical Analysis

#### 2.5.1. Behavior

Automatic scoring of total freezing and % freezing for each time point were performed by VideoFreeze software. Pre-tone, post-tone, and freezing measurements were calculated from the 20 s prior to and post tone and compared to the first 20 s of the tone presentation during each recall session.

#### 2.5.2. fUS

Temporal Power Doppler signal was extracted from ROIs defined via the Allen Mouse Brain Common Coordinate Framework (CCFv3) [35] individually aligned to each animal and imaging session (Fig. 2B) [34]. Power Doppler signal was processed using a similar method to [21,26,36], as follows. First, the signal was detrended with a 4^th^-order polynomial fit to remove potential low frequency drift. A 0.2 Hz low pass filter was then applied to isolate resting state functional connectivity frequencies [21,36]. Finally, Pearson product moment correlation coefficients were calculated for each ROI-ROI pair and normalized using Fisher’s *z* transformation to create a connectivity matrix. Node weight was calculated from the combination of all normalized correlation coefficients between a specific ROI and all other ROIs. To compare cue-evoked functional connectivity, Δ*z* values were calculated for each ROI-ROI pair by subtracting the connectivity value from the 60 seconds prior to the tone from that measured during the 60 second tone and averaged for the 3 tone blocks presented during each imaging session. Finally, those changes were compared across imaging sessions by calculating ΔΔ*z* values via the difference in these Δ*z* values.

#### 2.5.3. Statistics

All statistics were calculated using GraphPad Prism 8.4.3 software (California, USA). An ordinary one-way ANOVA with Dunnett’s multiple comparisons post-hoc test was used to compare freezing to tone to the pre-tone periods during fear expression, as well as to evaluate contextual freezing throughout the 5 minutes of exposure to the conditioning context. Significant differences in cue-evoked connectivity across imaging sessions were evaluated via a repeated measures two-way ANOVA with multiple comparisons and false discovery rate control by the Benjamini, Krieger, and Yekutieli two-stage step up method [37]. To investigate correlation between freezing behavior and functional connectivity, the Δ*z* of specific ROI-ROI pair clusters were compared to percent freezing using simple linear regression.

## 3. Results

### 3.1. Recent and remote responses to cued fear behavioral expression test

An associative memory trace formation was induced by training the animals with a Pavlovian cued fear conditioning (FC) task with a single tone – 1 mA foot-shock pairing. Recent memory retrieval was tested after 24 hours in a dissimilar context (context B), where fear conditioned animals displayed a marked increase in cue-associated freezing behavior (52.0 ± 9.4%, *p* < 0.001; Fig. 3). After 31 days, remote memory retrieval was tested. Here animals showed slightly reduced, but still significant, freezing behavior 36.9 ± 6.0% (*p* < 0.01), followed by an immediate resumption in motility. Fear conditioned animals displayed low levels of freezing (7.2 ± 2.7%) when re-exposed to the conditioning context A after 24 hours as well as minimal freezing behavior during the 1 minute pre-and post-tone time periods, demonstrating a tone-specific fear memory, as opposed to a context-associated memory. Three animals displayed less than 15% freezing after conditioning and were excluded from further analyses (FC, n=9) Animals that were only exposed to the unconditioned tone did not exhibit significant freezing behavior at any time (UT, n=12). Thus, the single, robust tone – shock pairing protocol was sufficient to induce a long-term Pavlovian fear memory.

**Fig. 3.**
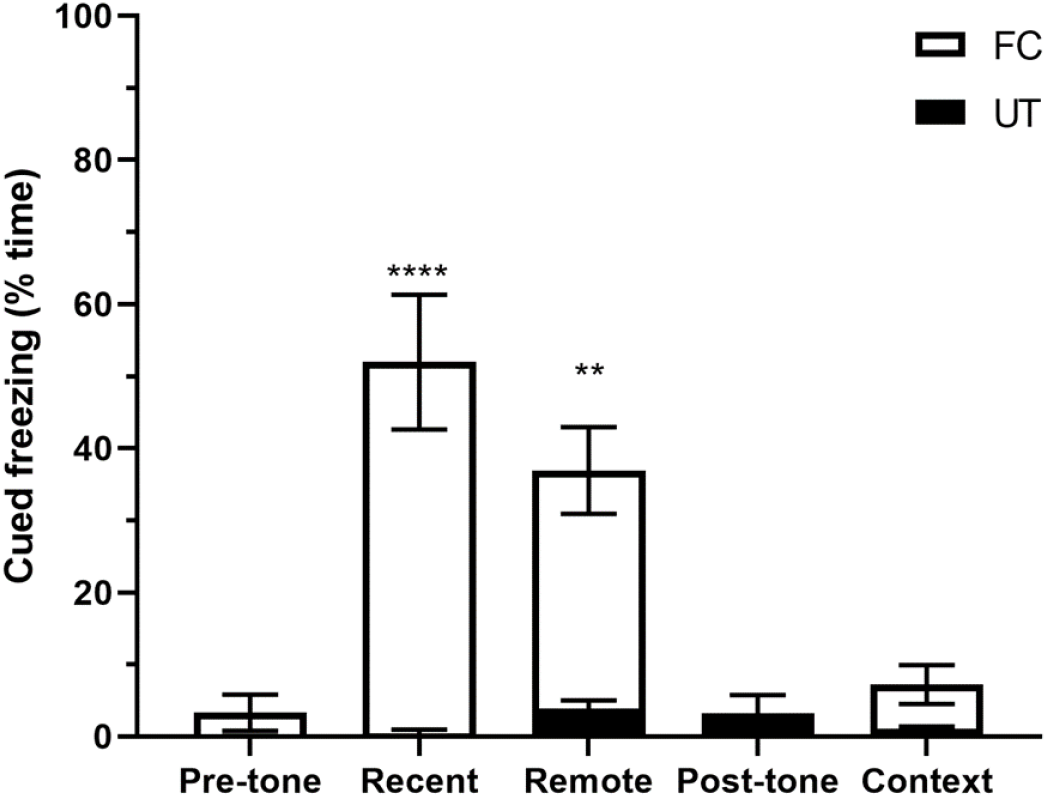
Average freezing responses to the auditory cue during recent and remote fear expression behavioral tests. FC animals showed significant freezing behavior during the tone in both recall sessions. UT trained animals showed negligible freezing during all time periods. When re-exposed to the conditioning context A after FC, animals exhibited minimal freezing (Context). Thus, validating the single robust US-CS pairing protocol for cued long term fear memory encoding. All data presented as mean ± SEM, significant differences between groups denoted with ^**^ or ^***^ (*p* <0.01, and *p* <0.001, respectively); (FC) fear conditioned group, n=9; (UT) unconditioned tone group, n=12.

### 3.2. Functional connectivity measurements

#### 3.2.1. Cue-evoked functional connectivity across recall imaging sessions

Imaging recall sessions were performed with functional ultrasound under light dexmedetomidine sedation. Cue-evoked changes were measured through the differences in functional connectivity prior to and during CS presentation (Δ*z*). A majority of ROI pairs showed increased connectivity during the CS in comparison to baseline in both the recent and remote recall sessions in fear conditioned animals (Fig. 4A). Scaled matrices show the differences between imaging sessions in both the FC and UT groups for all ROI pairs (ΔΔ*z*, Fig. 4B-D). Significant increases in functional connectivity of the LA with the PL, HC, BLA and CeA were apparent in both recall sessions in the FC group (Fig. 4B & C, lower triangles). During the recent recall session, increases in connectivity of the TeA with the A1, amygdalar nuclei, along with inter-connectivity among the CeA with the PL, HC and amygdalar nuclei were significant (Fig. 4B, lower triangle), whereas these connections were still increased from baseline (Fig. 4C, lower triangle), but reduced in the remote recall session in comparison to the recent session (Fig. 4D, lower triangle). Interactions with the A1, IL, BMA and HC were also increased during remote recall (Fig. 4C, lower triable), but showed both increases and decreases in comparison to the recent recall session (Fig. 4D, lower triangle). The UT group did not exhibit widespread changes in que-evoked connectivity (Fig. 4B-D, upper triangles). Overall connectivity was lower throughout the brain during the tone presentation during both the remote and recent recall sessions compared to baseline, especially between the LA and BLA/BMA although some slight increases were observed in PL-TeA and PL-BLA connectivity.

**Fig. 4.**
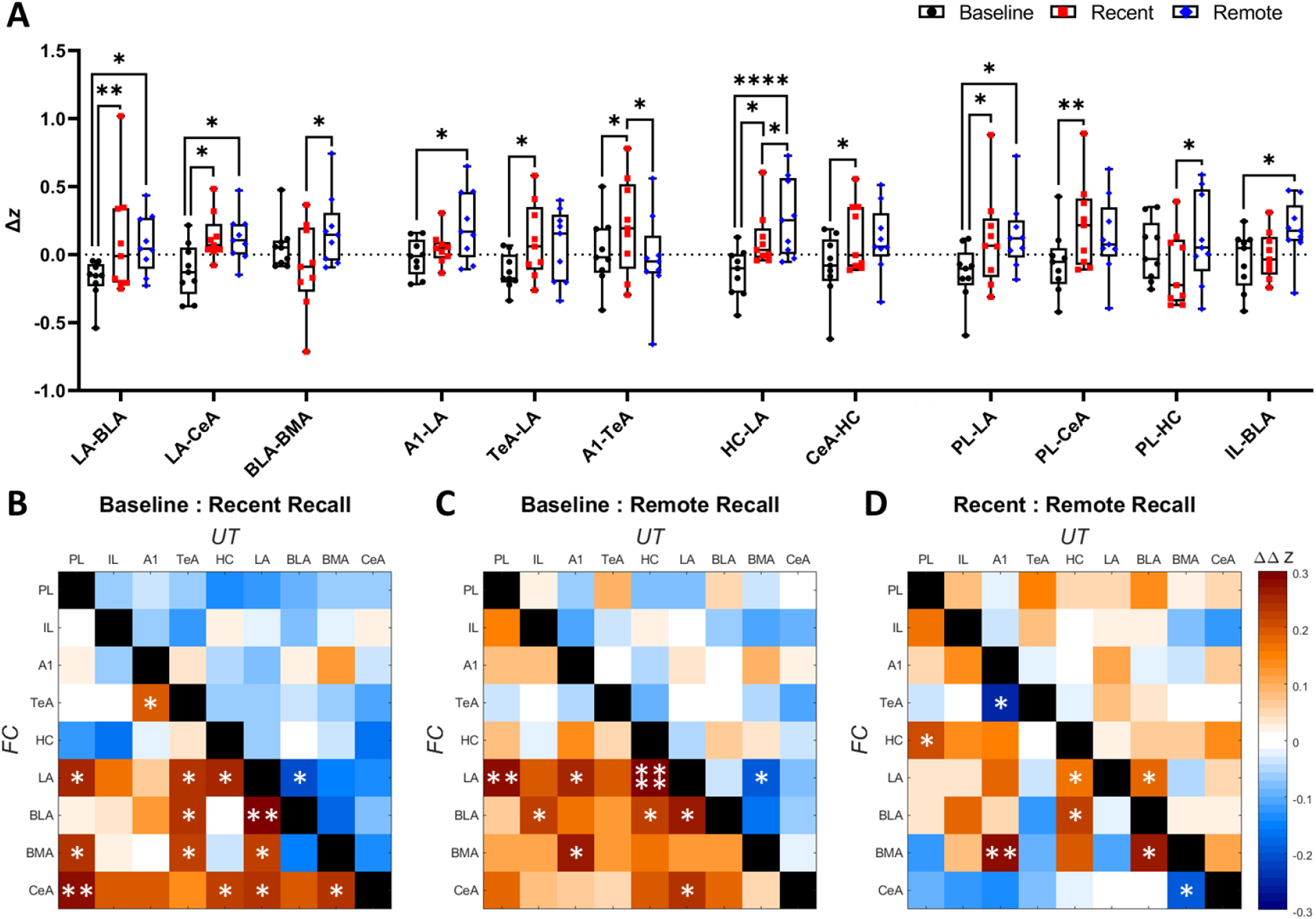
Functional connectivity changes in response to CS presentation between imaging sessions. (A) Intra-session changes in connectivity for select ROI-ROI pairs in FC group. Inter-session differences in tone-evoked connectivity between baseline and recent recall sessions (B), baseline and remote recall sessions (C), and recent and remote recall sessions (D). FC group data in lower triangles, n=9, UT group in upper triables, n=12. Data presented as median ± 10-90 quartile Δ*z* (A), and mean ΔΔ*z* (B-D). Significant differences between imaging sessions denoted with ^*, **^ or ^****^ (*p* <0.05, *p* <0.01, and *p* <0.0001, respectively). Abbreviations defined in figure 2.

#### 3.2.2. Functional connectivity brain networks

Significant functional brain network comparisons were depicted using spheres for nodes and the interconnecting lines as edges (Fig. 5). The inter-session differences of total correlation coefficients for each ROI were used to calculate the node diameter size. Edge width and shade represent inter-session differences for specific ROI-ROI pair connectivity (ΔΔ*z*) where the greatest increases are represented as wide dark lines and decreases as white dashed lines. Only edges passing hypothesis testing (Fig. 4B-D) are depicted. During recent recall the CeA, LA, and TeA nodes displayed the greatest increases in connectivity compared to baseline, as reflected by the increased sphere diameter and number of significant edges (Fig. 5A). Whereas the PL, IL, A1, HC, LA, and BLA all showed increased node weight during remote recall (Fig. 5B). When comparing the two recall sessions, substantial decreases in TeA and CeA node weight from recent to remote recall was apparent, as well as increases in the HC and BLA nodes (Fig. 5C).

**Fig. 5.**
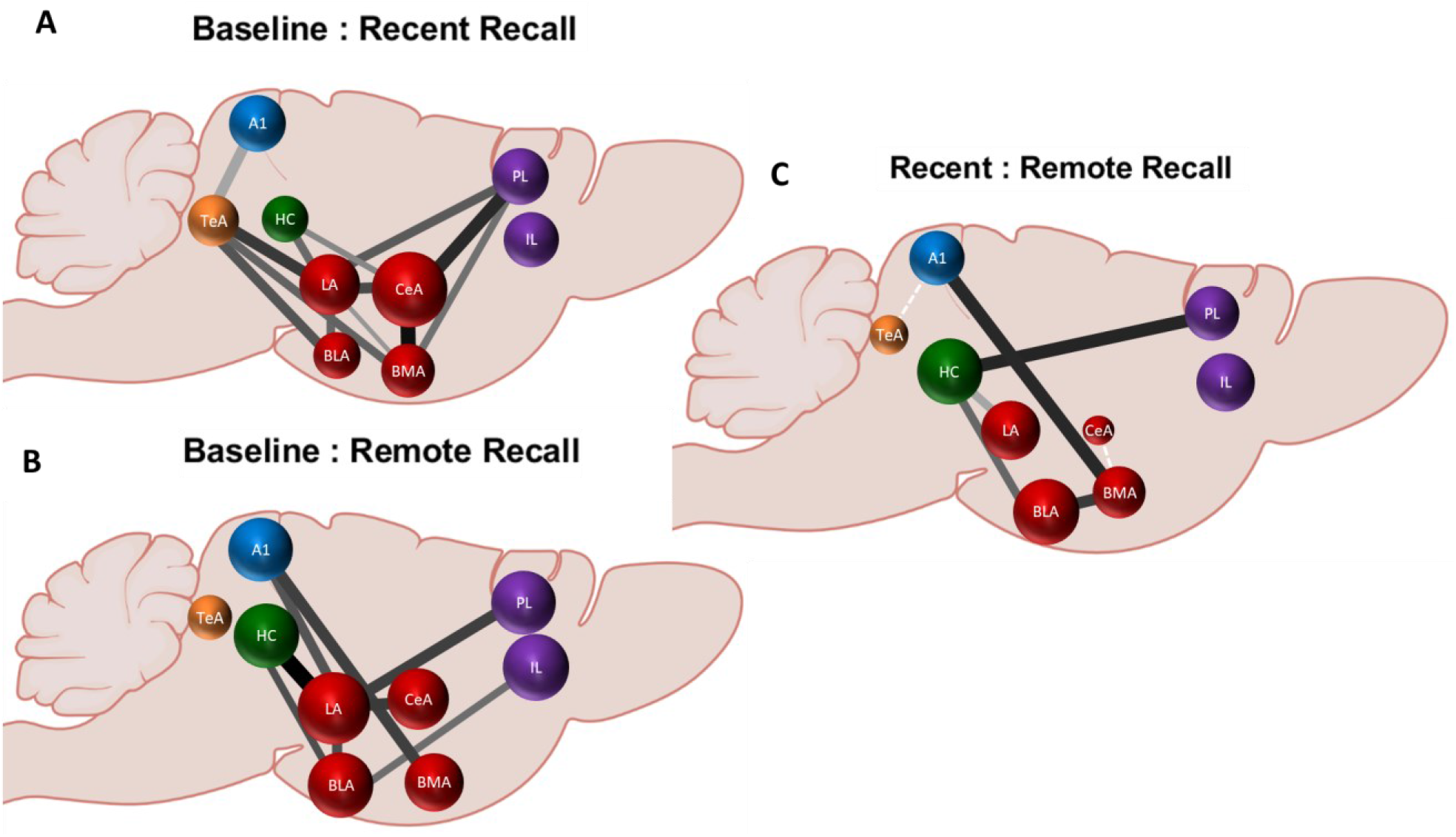
Brain network functional connectivity diagrams. The node diameter represents inter-session differences in global connectivity. Width and shade of edge color represent ROI-ROI functional connectivity changes (darker & wider = bigger increase), dashed white edges depict negative changes. Figure created with BioRender.com.

### 3.3. Correlation of freezing behavior with functional connectivity

Correlation between behavioral expression and functional connectivity showed positive and negative relationships depending on ROI-ROI pair cluster. Freezing behavior was positively correlated with PL, TeA, HC, and CeA functional connectivity during both recent and remote recall (Fig. 6A & C), while connectivity measures throughout the IL, A1 and BMA cluster were negatively associated (Fig. 6B & D).

**Fig. 6.**
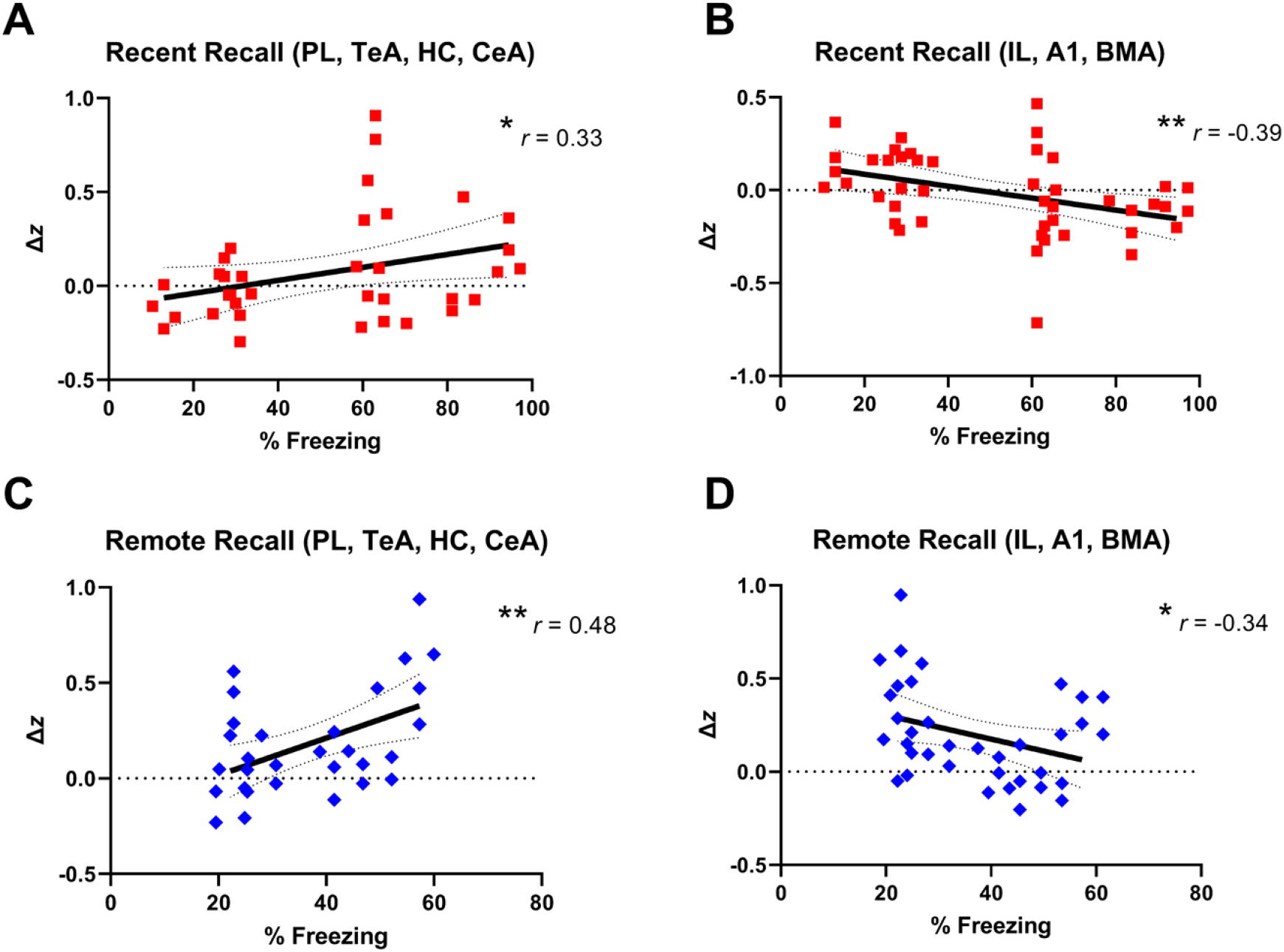
Correlation of behavioral fear expression and functional connectivity changes between clusters of ROI-ROI pairs in the FC group (n=9). (A,C) Connectivity between between PL, TeA, HC, and CeA nodes positively correlated with freezing behavior during both recent (A) and remote (C) recall sessions. IL, A1, and BMA inter-nodal connectiviy negatively correlated with freezing during recent (B) and remote recall (D). Each point represents a ROI-ROI pair within the denoted cluster for an individual animal. Significant correlations denoted with ^*^ or ^**^ (*p* < 0.05 and *p* < 0.01, respectively; simple linear regression; dotted line indicates 95% confidence interval).

## 4. Discussion

This study aimed to investigate the underlying neural functional connectivity in recent and remote fear memory retrieval in mice. In accordance with previous studies, one training session with a single tone-shock coupling induced the formation of a robust memory that could be detected during recent and remote recall sessions [31,32], one and thirty days after task acquisition, respectively (Fig. 3). In both memory retrieval sessions, robust cue-evoked increases in connectivity were detected throughout the associative fear network in the FC group, namely the LA, BLA, CeA, PL and HC (Fig. 4 & 5). In the remote recall session, however, the additional influence of neuroanatomical regions such as the IL and A1 along with reductions in global CeA and TeA cue-evoked connectivity were apparent (Fig. 4 & 5). Furthermore, regions associated with fear expression and memory consolidation (PL, TeA, HC, & CeA,) were positively correlated with freezing behavior during both recent and remote memory retrieval, while regions primarily associated with extinction learning (IL, A1, & BMA) were negatively associated (Fig. 6). In contrast, the UT control group did not display patterns of connectivity associated with fear memory. Thus, our data shows for the first time that recall of an associative fear memory trace can be detected with fUS in the lightly sedated mouse brain.

### 4.1. Increased intra-amygdalar connectivity is prominent during recent and remote retrieval memory traces

In both retrieval sessions FC mice showed significant tone-evoked increases in functional connectivity between numerous ROI-ROI pairs. Specifically, we found extensive increases in connectivity throughout the amygdala. The LA emerged as a major hub during both recent and remote recall sessions, where significant increases with the BLA, CeA, A1, HC and PL were detected (Fig. 4). These results correlate with existing literature that highlight the role of the LA as a convergence point for auditory and somatosensory inputs and a site for long-term potentiation associated gene induction, especially during an emotional memory [38,39]. The LA, however, is not sufficient for permanent memory encoding, rather a distributed amygdalar network is thought to be involved [6,11,40]. Intra-amygdalar connectivity between the LA and the BLA/BMA, together forming the basolateral complex, is well established and known to play a key role in emotion-related processes [10,41]. Together, the basolateral complex projects to the major output nuclei in the CeA [42]. Inter-connectivity between these regions was found to be significantly increased during CS presentation in both recent and remote fear memory recall sessions in FC mice, while the UT group displayed mainly insignificant decreases in connectivity (Fig. 4). Interestingly, while there was an increase in cue-evoked connectivity of the CeA with most ROIs during both retrieval sessions (Fig. 4B & C, bottom row of elements), the extent of this activation was decreased during the remote recall session (Fig. 4D, bottom row of elements). Concurrently, CeA node weight was also reduced in comparison to recent retrieval (Fig. 5C). In parallel, CeA connectivity was positively correlated with freezing behavior during both recall sessions (Fig. 6A & C). The underlying cause of this reduction could be diminishment of the memory trace engram, or alternatively the introduction of extinction behavior, a novel learning process where multiple presentations of the uncoupled conditioned stimulus results in a dampening of the fear response. These findings are comparable with those that have shown a fear extinction behavior-associated decrease in CeA output [41,43]. Cued fear extinction behavior can be detected after as few as two cue blocks [44–46], therein, this reduction of CeA connectivity during remote recall could be indicative of the start of extinction learning along with reflecting an overall dampening of fear expression.

### 4.2. Evidence of memory retrieval and consolidation in the HC and cortical sensory regions

The role of the hippocampus in cued fear memory retrieval is somewhat controversial. We detected elevated HC connectivity during both recall sessions and a large increase in node weight during the remote memory retrieval session (Fig. 4 & 5). Specifically, increased connectivity between the HC and LA was very prominent during both imaging sessions. Several studies have also detected a role for the HC in long term associative fear memory retrieval. Tone-induced increases in theta wave synchronicity, as well as auditory evoked potentials between the HC and LA, known to facilitate neuroplasticity, have been shown after successful FC [17,18]. It has been suggested that the HC is involved in reactivation of the associative fear memory trace, in compliment to its role in consolidation [19,47,48]. In the present study, however, it is difficult to determine if the HC is acting to retrieve an associative memory, is involved in a new learning process, or more likely a combination of these processes. The positive correlation between connectivity and freezing behavior (Fig. 6A & C) suggests that it is more likely that the HC is exhibiting greater involvement in the fear associated memory trace in the current study.

In addition, cue-evoked changes in the A1 and TeA also display signatures of memory consolidation. First, increased connectivity between the A1and TeA during recent memory retrieval demonstrates a classic signature of auditory processing, which was significantly reduced during remote recall. During remote recall, both the A1 and TeA showed cue-evoked increases in connectivity with the LA, in contrast to significantly lower inter-connectivity (Fig. 4). The TeA-A1 circuit has shown involvement in the recognition, discrimination, and encoding of auditory signals [49,50], important processes during recent associative memory retrieval. Our data shows that during remote recall, however, this intra A1-TeA cue-evoked connectivity increase is no longer present and both cortical regions exhibit stronger direct connections with the amygdala. These dynamic results suggest that during remote retrieval the memory engram cells in these intra-auditory areas are involved in the memory trace directly without inter-regional processing that is normally present during auditory memory encoding, a sign of memory consolidation.

### 4.3. PFC-amygdala connectivity is refined during remote fear memory retrieval

While there were dramatic changes in connectivity in the amygdala during both recall sessions, the PFC is thought to play a major, although dual, role in long-term memory retention [2,51]. Indeed, significant increases in functional connectivity were detected between the PL and amygdala, especially between the PL-LA, during both retrieval sessions. A widespread network between the PL and LA, BMA and CeA was apparent during the recent recall session (Fig. 5A). Although there was a reduction in connectivity between the PL and the more ventral amygdalar nuclei (Fig. 5B), there were increases in connectivity between the PL-LA as well as between the PL-HC from the recent to remote recall sessions (Fig. 4D). Refinement of PL connectivity has been shown after memory consolidation [52], which could suggest a shift from harmonizing the memory trace to a more coordinated and directive role in retrieval. In contrast, we saw an increase in IL connectivity to the BLA during remote recall session, which was absent during recent retrieval. The IL has been consistently shown to be a mediator of extinction learning [53–55]. The IL-BLA connection, in particular, has been frequently found to be crucial in extinction learning. An increase in intrinsic activity in the projection neurons between the two regions is found after extinction training and a reduction of this connectivity leads to failure of extinction memory retrieval [53,56]. Additionally, our finding that connectivity of the IL was inversely correlated with freezing behavior, suggests an active contribution of the IL and provides additional evidence for the presence of a fear extinction network.

### 4.4. Shock intensity

The present study used a single CS-US pairing with high shock intensity (1 mA), as this has previously been shown effective in maintaining a freezing response for several weeks after FC [32]. Yet, the question remains as to whether higher levels of freezing are indicative of a stronger fear association memory, or rather reflect an increased sensitivity to input received in close temporal proximity to the administration of the shock, given its high emotional impact [31]. However, the strong evidence from the correlation of functional connectivity with behavior, specifically the positive associations which were stronger during remote recall (Fig. 6A & C), suggest a relationship between behavior, connectivity, and the robustness of the memory. Another aspect to consider is the conditioning protocol used in the current study. Here, a single intense shock (1 mA, 2 s) was delivered, whereas other protocols using multiple presentations of lower intensity shocks are also common (i.e., 3 × 0.2 to 0.6 mA) [57,58]. Increasing the number of CS-US pairings could induce an alternative memory trace where the cortical involvement is stronger, thereby shifting the core of connectivity increases from the amygdala to the cortex, especially during remote memory retrieval. However, care must be taken to ensure that the fear memory remains specific to the auditory cue and not coupled to the conditioning context.

### 4.5. Implications

The ability to acquire and analyze circuit level data is crucial when investigating a complicated behavior. Animal models of human behavior are a valuable tool for the in-depth study and manipulation of specific cell types, brain regions, and behaviors and their impact on neural networks. Techniques such as fMRI and fUS take advantage of blood flow changes as an indirect measure of neural activity to image the brain on a large scale. While most human circuitry data comes from fMRI, this technique is less suitable for small animals due to the need for a stronger magnetic field for higher spatial resolution, which typically comes at a cost of temporal resolution. In addition, the loud vibration of the gradient coils characteristic in MRI can reach 80 – 115 dB [59], limiting the applicability of auditory stimuli and adding the complication of training for awake or lightly sedated animals. Therefore, fUS could fill this translational gap and provide a tool for both forward and back translation, drug development, and behavioral interventions in numerous psychological disorders. fUS has been more recently applied in non-human primates (NHPs) and in some clinical applications as well. The ability to use the same technique in small animals, where a whole network can be visualized, and then upscale these results to NHPs and humans opens a broad range of applications for fUS and network analysis. Indeed, fUS has recently been applied in the decoding of movement intentions which could have a major role in less invasive brain machine interfaces for a wide variety of disorders [22,60–62].

### 4.6. Limitations and future considerations

While it would be ideal to image awake and behaving animals with fUS, an advancement currently in development [36,63–65], intact functional connectivity and stimulus responsiveness have been shown using similar dexmedetomidine sedation doses to those used here [26,29]. Furthermore, the effects of anesthesia and heavy sedation have been shown to reduce connectivity, lending more weight to the significance of our results. Nonetheless, the capability to image animals during FC acquisition would open a new line of study and could shed light into the phenomenon of memory formation, an endeavor that would be logistically and technically very difficult with other imaging modalities. Also, we used a linear array ultrasonic probe with a dedicated global positioning system [34] to isolate a single cross-sectional imaging plane; volumetric coverage of the whole brain would allow empirical evaluation of a more distributed network. The development of matrix arrays will overcome this limitation, although they currently suffer a sacrifice in spatial resolution [66,67]. While the current linear probe offers excellent spatial and temporal resolution compared to most imaging modalities, an increased rate of compiled images would be needed to determine directionality of functional connectivity. This has been accomplished using a probe with more channels, reducing the number of compounded images, and employing a sliding window data analytics approach where a directional time delay as short as 0.27 s between regions has been measured [62]. Employing this approach would enable isolation of top-down and bottom-up pathways of fear memory retrieval. Finally, although we found evidence for the introduction of extinction learning, we did not explicitly employ an extinction learning paradigm protocol. It would be of interest to explore this facet in future studies to evaluate whether extinction learning shifts the balance between these PL - amygdala and IL - amygdala memory traces, or if the associative fear memory is overwritten. Employing a protocol with multiple low-intensity US shocks could also be explored to investigate a potential shift of the core of the memory trace from the amygdala to more cortical regions.

## 5. Conclusions

The present study showed for the first time that the neural circuitry underlying emotional memory encoding and long-term storage can be investigated in a preclinical animal model with functional ultrasound, a state-of-the-art imaging technique. Similar to previous studies, the lateral amygdala and central amygdalar nuclei emerged as hubs of cue-evoked functional connectivity. We also found evidence for hippocampal involvement in recent and remote memory retrieval. Both the prelimbic and infralimbic cortices showed variable changes in connectivity, with a refinement in connections evident during remote recall. Regions known to be relevant in the fear association network positively correlated with freezing behavior, while regions associated with extinction behavior were negatively associated, giving credence to the suggestion that strength of functional connectivity underlies the strength of the memory. Thus, we showed for the first time that fUS is a powerful tool for elucidating large scale neural networks underlying associative fear memory formation and storage, providing insight to guide further studies in the development of novel treatments for anxiety related and affective disorders.

## CRediT authorship contribution statement

GGM: Conceptualization, Methodology, Investigation, Visualization, Formal Analysis, Writing – Original draft, Writing – Review & Editing, Project Administration; RS: Conceptualization, Investigation, Formal Analysis, Writing – Original draft, Writing – Review & Editing; JG: Writing – Review & Editing; TB: Supervision, Writing – Review & Editing; BH: Funding acquisition, Conceptualization, Supervision, Writing – Review & Editing.

## Acknowledgements

The authors would like to thank Chris Pryce, Hannes Sigrist, Serena Deiana, and Carsten Wotjak for their expertise and advise in the planning of fear conditioning experiments. We would also like to thank Jeremy Ferrier, Artem Shatillo, and Eliane Proulx for their advice in fUS interpretation and analytical techniques. We also thank Ester Nespoli for her recommendations in review of this manuscript.

